# D- and L-Lactate enhance intestinal barrier function via activation of an apical HCAR1/Gαi pathway in a human colonic epithelial cell model

**DOI:** 10.1101/2025.10.28.685091

**Authors:** Annabelle J. Milner, Priyanka Anujan, Gary S. Frost, Aylin C. Hanyaloglu

## Abstract

The stereoisomers of lactate, L- and D- are not only metabolic substrates but also signalling molecules, capable of activating and signalling through its G protein-coupled receptor, Hydroxycarboxylic acid receptor 1 (HCAR1). These stereoisomers are both produced by the gut microbiota at millimolar concentrations creating a physiological environment for lactate-sensing unique to the gut yet, poorly understood. Here we identify a role for D-/L-lactate on intestinal barrier function. A human colonic epithelial cell model, Caco2, activated Gαi signalling in response to both L- and D-lactate, although L-lactate exhibited a more potent and rapid Gi signal profile. When differentiated, apically but not basally treated D-/L-lactate enhanced tight junctions and reduced cell permeability, consistent with the apical localization of HCAR1. This improved barrier function occurred in a Gαi-dependent manner. In addition, apical lactate rescued the reduced intestinal barrier function induced by lipopolysaccharides. This work highlights the potential for D-/L-lactate supplementation in improving gut health.

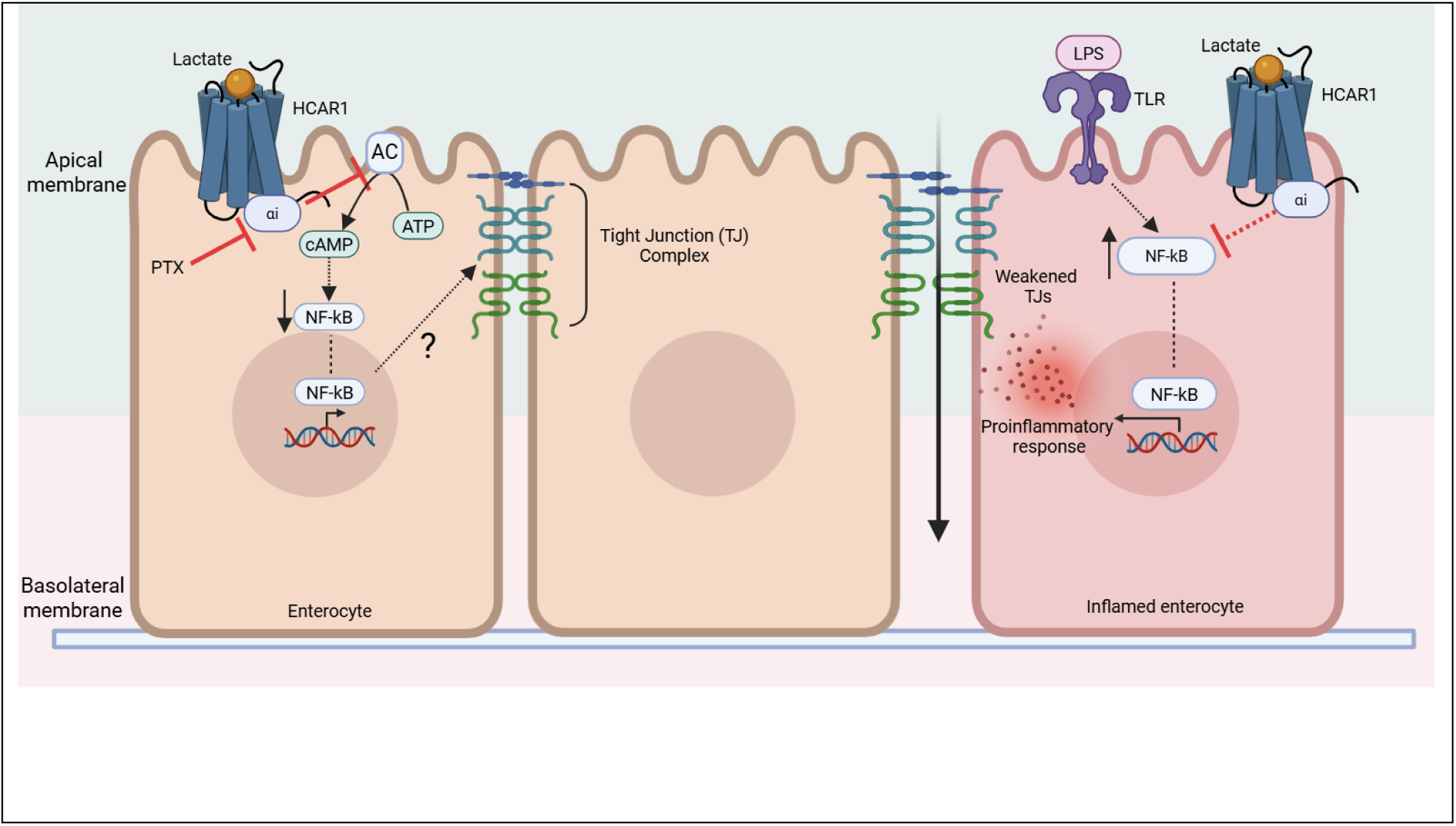

## Introduction

The original concept of lactate acting as just a metabolic waste product has undergone a profound transformation over the past three decades. It is now recognised as a multifunctional molecule, serving as an energy substrate, a metabolite for gluconeogenesis, and mediates cell signalling through G-protein Coupled Receptor (GPCR) activation (Brooks, 2021). Advances in selective analytical techniques have further enabled for the differentiation between the stereoisomers of lactate: L- and D-lactate (Pohanka, 2020). While the physical structure of these enantiomers is similar; the synthesis and metabolism differ vastly. L-lactate can be synthesised via one of two mechanisms: glycolysis or bacterial fermentation, whereas D-lactate is primarily only formed through bacterial fermentation by the gut microbiota (Rao et al., 2018). Consequently, this makes the colon one of the only places in the human body where both L- and D-lactate are synthesised in sufficient quantities to elicit biological effects under non-pathophysiological conditions.

The interaction between metabolites produced by the gut microbiota and intestinal epithelial cells has long been of interest due to its influence on local signalling pathways, and its broader impact on gut health (Ghosh et al., 2021). These microbial-derived metabolites not only directly interact with intestinal epithelial cells but also peripheral immune cells to modulate immune activities. Intestinal epithelial cells form a physical and biochemical barrier that facilitates nutrient absorption while preventing the passage of harmful substances and pathogens. Tight junction (TJ) complexes comprising of transmembrane (Claudins, Occludin, and Junctional Adhesion Molecules), and cytoplasmic Zonula Occluden (ZO) proteins provide structural support for maintaining close cell-cell contact, thus forming a tight, selective barrier. TJs additionally regulate the paracellular and transcellular transport of substances across the epithelial barrier. Disruptions in TJ integrity and/or intestinal epithelial barrier function have been associated with ‘leaky gut syndrome’, a precursor to numerous pathogenic diseases including intestinal bowel disease and celiac disease. While it remains debated whether barrier dysfunction precedes or results from disease onset, evidence suggests that dysbiosis-induced increased barrier permeability supports the former hypothesis (Horowitz et al., 2023).

Lactate has been identified as the endogenous ligand for GPR81, also known as Hydroxycarboxylic acid receptor 1 (HCAR1). A member of the Class A GPCR family, HCAR1 couples to the Gαi/o family of heterotrimeric G proteins to inhibit adenylate cyclase activity and decrease levels of the intracellular second messenger cyclic AMP (cAMP). This HCAR1-Gαi/o signal pathway has been implicated in modulation of lipolysis, inflammation, and cancer progression (Liu et al., 2009; Feingold et al., 2011; Feng et al., 2017). Local microbiota-derived lactate concentrations in the gut can exceed 10 mM, compared to circulating blood lactate levels of 0.5-2 mM, prompting growing interest in the role of lactate-mediated HCAR1 signalling in intestinal epithelial function. Lactate signalling via HCAR1 has been shown to stimulate intestinal stem cell proliferation and renewal, emphasising the receptor’s importance in maintaining gut homeostasis (Lee et al., 2018). Additional studies have demonstrated that mouse HCAR1 knockout increases susceptibility to colonic inflammation (Ranganathan et al., 2018) and that lactate provides protection against experimental-induced colitis (Iraporda et al., 2016;Li et al., 2024).

While these studies underscore the beneficial effects of lactate on intestinal homeostasis, the underlying molecular mechanisms remain poorly understood, with most research being conducted in mouse models. To address this gap, we investigated L- and D-lactate-induced signalling and their roles in modulating intestinal barrier integrity using the human colonic epithelial Caco-2 cell line, which endogenously expresses HCAR1, including a line generated that expresses epitope-tagged HCAR1. Once differentiated, Caco-2 cells exhibit apical-basal polarity enabling simulation of both luminal and blood-derived basal lactate exposure, therefore we assessed D-/L-lactate induced HCAR1 signalling and trafficking and the effects of Caco-2 cells to sense lactate sidedness. We identified that apical HCAR1/Gαi activation enhances barrier integrity and reverses inflammation-induced damage to colonic epithelial cell barrier integrity.

## Results

### L- and D-lactate induce distinct cAMP activation profiles in Caco-2 cells

To investigate the role of lactate in maintaining intestinal homeostasis, a human colonic epithelial cell line, Caco-2, was employed. HCAR1, the only known GPCR to be activated by lactate, was not only endogenously expressed in Caco-2 cells (Human Protein Atlas and Supplemental Figure 1A) but could also be activated by lactate at physiologically relevant concentrations (Figure 1). Caco-2 cells exposed to either L- or D-lactate inhibited forskolin-induced intracellular cAMP accumulation in a concentration-dependent manner (Figure 1A). EC_50_ values were determined to be 1.30 ± 0.25 mM and 2.90 ± 0.12 mM (mean ± SD) for L- and D-lactate, respectively indicating that L-lactate was significantly more potent than D-lactate. The use of a Gαi inhibitor, pertussis toxin (PTX), confirmed that both L- and D-lactate stimulated forskolin-induced reduction in cAMP levels was dependent on Gαi activation (Figure 1A). Kinetics of Gαi signalling, assessed using the real time GloSensor cAMP sensor, for L- and D-lactate confirmed differences between the stereoisomers in half-maximal inhibitory profiles (Figure 1C-E). L-lactate elicited a faster Gαi signalling response compared to D-lactate, with half maximal inhibition of 152.00 ± 7.21 sec (L-Lactate, 10mM) compared to 239.33 ± 17.93 seconds (mean ± SD, D-lactate, 10mM) (Figure 1B-D).

**Figure 1.**
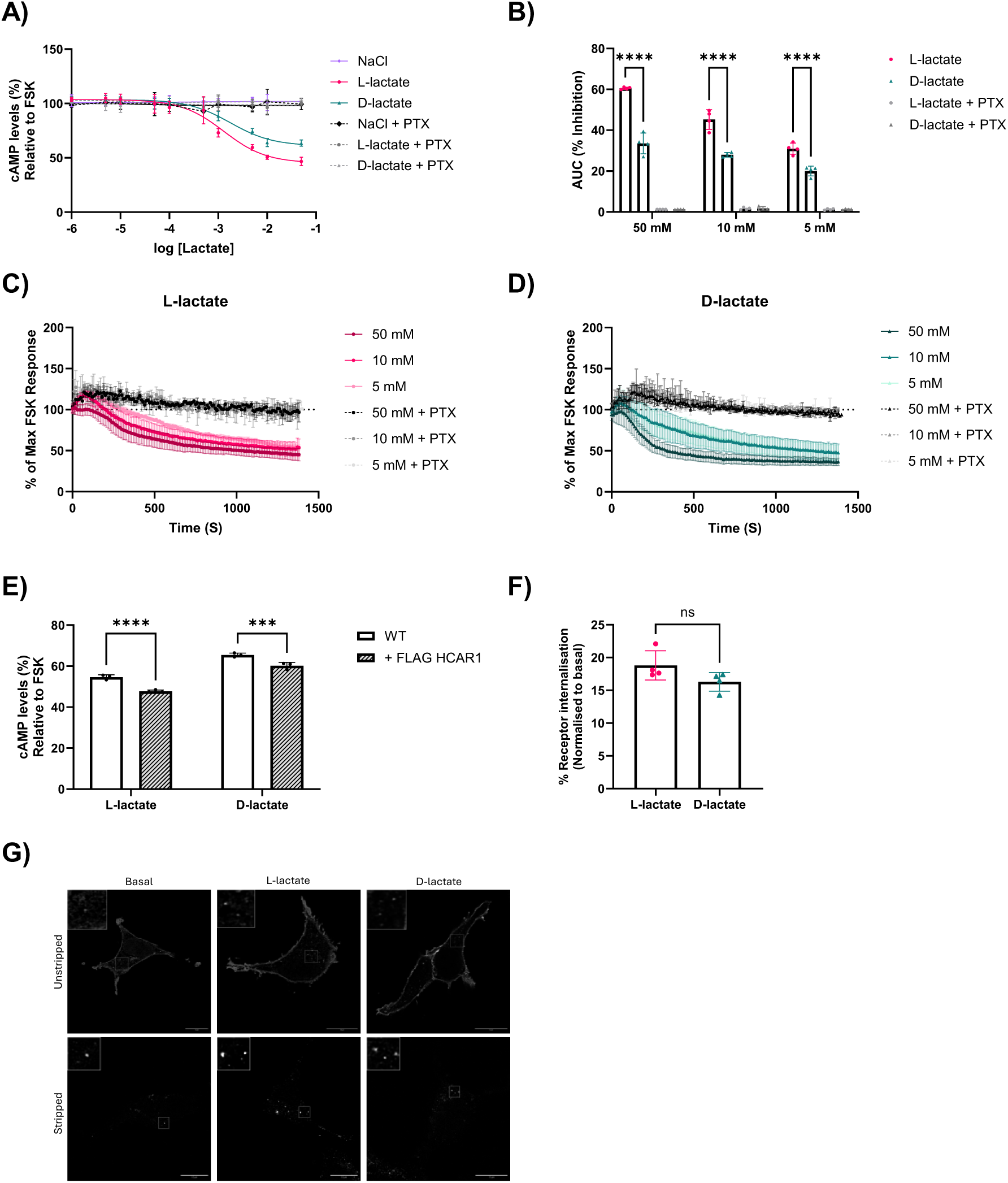
Both L- and D-lactate activate Gαi/o signalling in Caco-2 cells. A) Intracellular accumulation of cAMP levels measured in Caco-2 cells. Cells were pre-treated with or without PTX for 16 h, followed by IBMX (0.5 mM, 5 minutes), and then forskolin (3 µM), and NaCl, L- or D-lactate (100 mM-10 µM, 5 minutes). cAMP concentrations were normalised to protein and presented as percentage of maximal FSK-induced cAMP accumulation (100 %). Data presented as mean ± SEM, N=3. B-D) Caco-2 cells were transfected with 100 ng of Glo-sensor plasmid 24 h prior to ligand stimulation and pre-treated with or without PTX for 16 h, followed by forskolin (3 µM) and C) L-lactate, or D) D-lactate (50 mM, 10 mM or 5 mM). Luminescence levels measured for 25 minutes and traces plotted. B) AUC was presented as the net between forskolin AUC and lactate AUC. Data presented as mean ± SEM, N=4. One-way ANOVA, with Tukey’s post-hoc test, ***P<0.005, ****P<0.0001. E) Intracellular accumulation of cAMP levels measured in WT Caco-2 cells and Caco-2 FLAG-HCAR1 cells. Cells were stimulated with IBMX (0.5 mM, 5 minutes), and then forskolin (3 µM) and L- or D-lactate (10 mM, 5 minutes). Data presented as mean ± SEM, N=3. One-way ANOVA, with Tukey’s post-hoc test, ***P<0.005, ****P<0.0001. F) Cell surface receptor expression was measured in Caco-2 cells FLAG-HCAR1 cells following L- or D-lactate (10 mM, 20 minutes) stimulation. Data calculated as mean fluorescent intensity multiplied by % gated value, normalised to basal conditions (100 %). Difference between basal and ligand-induced conditions plotted as receptor internalisation. Data presented as mean ± SEM, N=4. Unpaired two-tailed Student’s T-test, Ns; not significant. G) Representative confocal images of Caco-2 FLAG-HCAR1 cells that were fed live labelling of anti-FLAG M1 antibody for 20 minutes followed by L- or D-lactate stimulation (10 mM, 20 mins) and fixed. Stripped condition were washed with EDTA+ to remove plasma membrane staining. Insert image highlights region of interest and is scaled up by 3. Scale bar 10 µM.

It is well documented that GPCRs can activate extracellular signal-regulated protein kinase (ERK1/2) signalling, and therefore the effects of L- and D-lactate stimulation on the phosphorylation levels of p44/42 ERK1/2 were assessed. Both L- and D-lactate stimulation resulted in peak p-ERK activity at 5 minutes before returning to basal. Caco-2 cells treated with PTX and then stimulated with L-lactate resulted in little ERK activation, suggesting that the response observed was due to Gαi signalling (Supplemental Figure 1B-C). In contrast, PTX-treatment followed by D-lactate stimulation did not fully suppress ERK activation, suggesting that D-lactate may subsequently increase ERK activation via a Gαi-independent mechanism (Supplemental Figure 1B-C), highlighting differences in how the stereoisomers may regulate HCAR1 signalling.

To employ a cell system where HCAR1 protein could be visualised, and due to a lack of commercially available selective and sensitive anti-HCAR1 antibodies, we generated an N-terminally FLAG-tagged HCAR1 that was stably expressed in Caco-2 cells and retained FLAG-HCAR1 expression through differentiation. cAMP profiles in response to L- or D-lactate (10 mM) were determined in undifferentiated Caco-2 FLAG-HCAR1 cells. The stable cell line exhibited a greater response to either L- or D-lactate-dependent inhibition on forskolin-induced cAMP levels compared to WT Caco-2 cells, (Figure 1E), which is likely due to increased receptor levels. Internalisation profiles of HCAR1 in response to lactate stimulation were determined qualitatively through confocal microscopy and quantified via flow cytometry. Caco-2 FLAG-HCAR1 cells were fed anti-FLAG M1 antibody live followed by L- or D-lactate stimulation (10 mM). A subset of cells was then washed with EDTA to remove plasma membrane receptor bound FLAG antibody (stripping). Confocal imaging revealed that the receptor is predominantly at the plasma membrane, with little to no constitutive internalisation in basal conditions (Figure 1G). Both L- and D-lactate promoted low levels of receptor internalisation, which can be observed in ‘stripped’ conditions. Quantification of receptor internalisation via flow cytometry demonstrated L-lactate promoted ∼18 % decrease in receptor at the plasma membrane, whereas D-lactate promoted ∼16 %, in comparison to basal conditions (Figure 1F).

### HCAR1 expression increases during Caco-2 cell differentiation

Once differentiated, Caco-2 cells resemble intestinal epithelial cells and express accordant characteristics such as microvilli and tight junctions (Lea, 2015). Here, we evaluated monolayer integrity and enterocyte morphology through transepithelial electrical resistance (TEER) readings, biochemical analysis and confocal microscopy across a 21-day differentiation period. In agreement with prior studies (Johannessen et al., 2013; Srinivasan et al., 2015; Yang et al., 2018), TEER readings increased and began stabilising around day 14 before plateauing (Figure 2B). Coinciding with this, expression of tight junction proteins ZO-1, Occludin, and Claudin-1 increased over the 21-day period. Furthermore, Ezrin and IAP, proteins used to assess microvilli formation and function, also increased by day 18 (Figure 2C-E). Enterocyte markers routinely used to identify cell populations within single-cell RNAseq datasets such as SLC26A2, CA2 and FABP1, were assessed in differentiating Caco-2 cells, however, expression levels of these proteins did not increase during differentiation (Supplemental Figure 2). On the contrary, mRNA levels of HCAR1 significantly increased by day 18 of differentiation (Figure 2F) suggesting that differentiation-induced polarity may be important in the regulation of HCAR1 expression.

**Figure 2.**
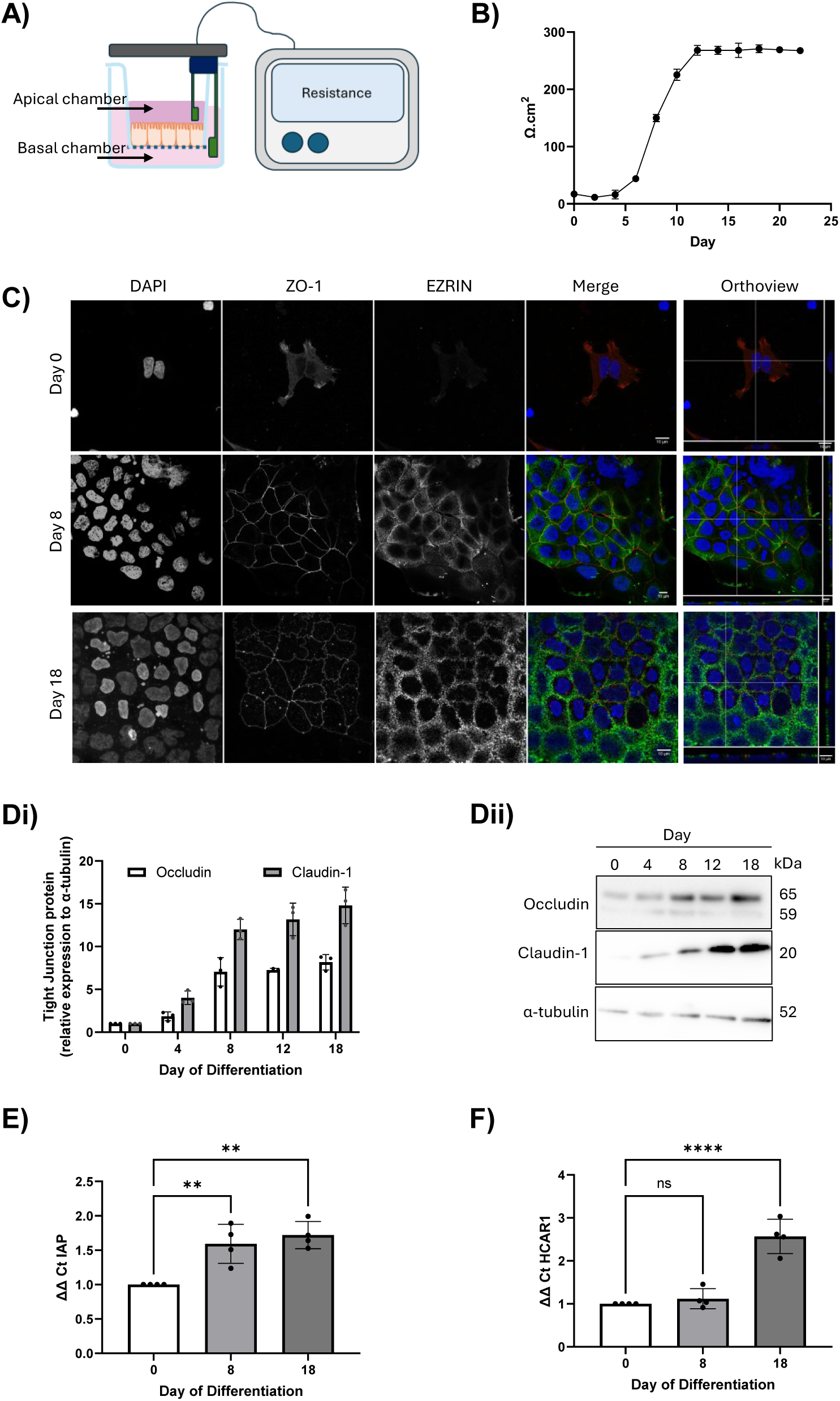
Characterising Caco-2 cell differentiation. A) Schematic showing measurement of transepithelial electrical resistance (TEER) readings in a Transwell system. B) TEER values of Caco-2 cells during differentiation. Caco-2 cells were grown on Transwell membranes for 22 days with apical and basal media changes and TEER readings every two days. Blank resistance was subtracted from TEER readings before being corrected for surface area of Transwell. Three TEER readings were taken per well and experiments ran in duplicates. Data presented as mean ± SEM, N=3. C) Representative confocal images of Caco-2 cells during differentiation. On day 0, 8 or 18, cells were washed, fixed, permeabilised and labelled for ZO-1 and ezrin. Cells were then labelled with immunofluorescent secondary antibodies AlexaFluor 488 (ezrin) and AlexaFluor 647 (ZO-1). Full Z-stacks of cells taken; images are max intensity of compressed Z-stack. Scale bar 10 μM. Orthoview of XZ and YZ profiles of differentiating Caco-2 cells. Scale bar 10 μM. D) Occludin and Claudin-1 protein expression levels analysed by western blot, measured in differentiating Caco-2 cells, and Dii) representative western blot. Expression normalised to α-tubulin. Data presented as mean ± SEM, N=3. E) IAP, and F) HCAR1 mRNA levels measured in differentiating Caco-2 cells. Relative abundance was analysed using ΔΔCT method, relative to GAPDH. Data presented as mean ± SEM, N=4. One-way ANOVA followed by Dunnett’s multiple comparison post-hoc test.

### HCAR1 is apically localised and its overexpression increases cell barrier integrity

Protein markers were used to confirm polarisation orientation of differentiated Caco-2 cells; Ezrin, a protein expressed within microvilli structures, was used as an upper apical marker, ZO-1, a protein expressed in tight junctions, as an apico-lateral marker, and a nuclear stain was used to define the basal position of the cell (Figure 3A). These markers enabled the generation of spatial fluorescence intensity plots, confirming that Caco-2 cells exhibited apicobasal polarisation, a hallmark of epithelial cells (Figure 3B-D). From here, using the spatially controlled protein markers as orientational references, the cellular localisation of FLAG-tagged HCAR1 stably expressed in Caco-2 cells could be determined. Analysis of fluorescence intensity plots indicated that HCAR1 was primarily expressed on the apical surface of the polarised intestinal epithelial cell (Figure 3E-G).

**Figure 3.**
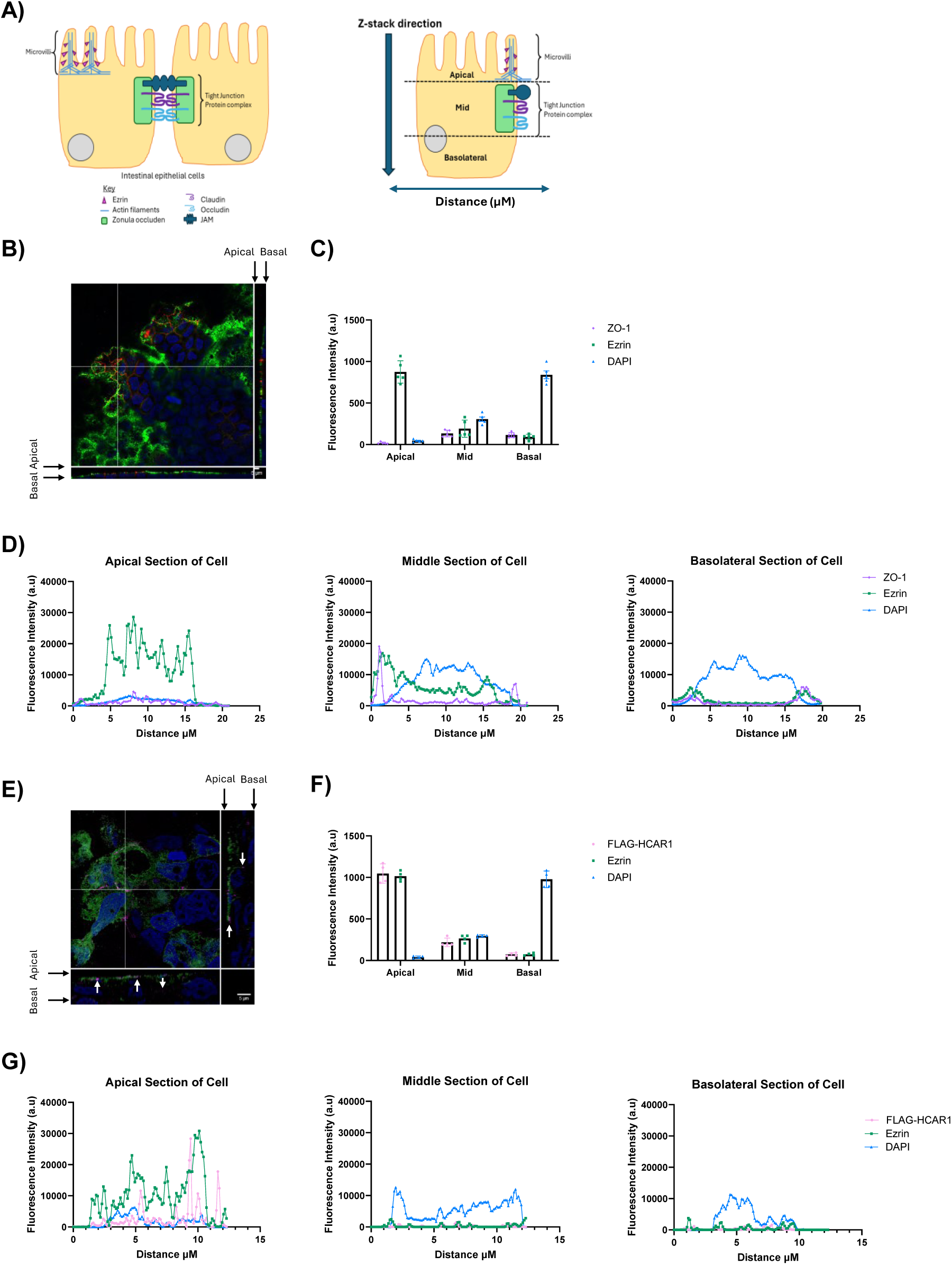
Characterisation of the spatial expression of FLAG-HCAR1 in differentiated Caco-2 cells. A) Schematic showing markers of polarisation in Caco-2 cells and Z-stack orientation. B) WT Caco-2 cells were differentiated for 18 days and washed, fixed and permeabilised for either ezrin, ZO-1, or DAPI. Cells were then labelled with immunofluorescent secondary antibodies AlexaFluor 488 (ezrin-green) and AlexaFluor 647 (ZO-1-purple). Full Z-stacks of cells taken; Representative Orthoview of XZ and YZ profiles of differentiated Caco-2 cells. C) Fluorescence intensity plots of each protein expression in the upper third section of the cell termed apical, middle third, termed middle and lower third section termed basolateral. Fluorescent intensity was determined using the multiple line intensity profile plot tool in ImageJ. Data presented as mean ± SEM, N=4 where three individual cells were quantified for each biological repeat. D) Representative fluorescent intensity profiles of ezrin, ZO-1, and DAPI at select Z-slices of a differentiated WT Caco-2 cell. Confocal images were acquired using the same gain and intensity settings and full Z-stacks were captured. E) Representative Orthoview of XZ and YZ profiles of Caco-2 FLAG-HCAR1 cell stained for FLAG-HCAR1 (pink), ezrin (green) or DAPI (blue). White arrows point towards FLAG-HCAR1. Scale bar 5 μM. F) Fluorescence intensity plots of HCAR1, ezrin and DAPI in the apical, middle and basolateral section of Caco-2-FLAG-HCAR1 cells. Data presented as mean ± SEM, N=3 where three individual cells were quantified for each biological repeat. G) Representative fluorescent intensity profiles of FLAG-HCAR1, ezrin and DAPI at select Z-slices of a differentiated Caco-2-FLAG-HCAR1 cell. Confocal images were acquired using the same gain and intensity settings and full Z-stacks were captured.

The morphology and characteristic behaviour of Caco-2 FLAG-HCAR1 cells were comparable to WT Caco-2 cells, however, while differentiating into polarised epithelial cells, the TEER readings were elevated in the Caco-2 FLAG-HCAR1 cells (Figure 4A). Additionally, TEER readings in Caco-2 FLAG-HCAR1 cells plateaued earlier, potentially suggesting a role of HCAR1 in intestinal barrier integrity. To examine if increases in TEER readings in the FLAG-HCAR1 cells corresponded to changes in cell permeability, FITC Dextran (4 kDa) was employed. FITC Dextran (1 mg/mL) was added to the apical chamber of the transwell and sample measurements were collected from the basolateral chamber across different time points. Equivalent media was added to the basolateral chamber to prevent bulk fluid flow imbalance. The apparent permeability (P_app_) for WT Caco-2 cells in basal conditions was calculated to be 3.91 × 10^7^ cm.s^-1^ whereas, the P_app_ for Caco-2 FLAG-HCAR1 cells was determined to be 6.13 × 10^8^ cm.s^-1^ (Figure 4B), therefore, not only did differentiated FLAG-HCAR1 cells have tighter forming tight junctions, but equally, a lower paracellular flux. As lactate can be sensed by monocarboxylic acid transporter 1 (MCT1) and sodium-monocarboxylic acid transporter (SMCT) 1 and 2, expression levels of these transporters were assessed during differentiation of Caco-2 and Caco-2 FLAG-HCAR1 cells to ascertain if the enhanced effects on increased TEER readings in this cell line could be attributed to lactate acting through both HCAR1 and the transporters. Expression levels of MCT1, SMCT1, and SMCT2 were consistent throughout differentiation and were comparable between the two cell lines (Supplemental Figure 3A).

**Figure 4.**
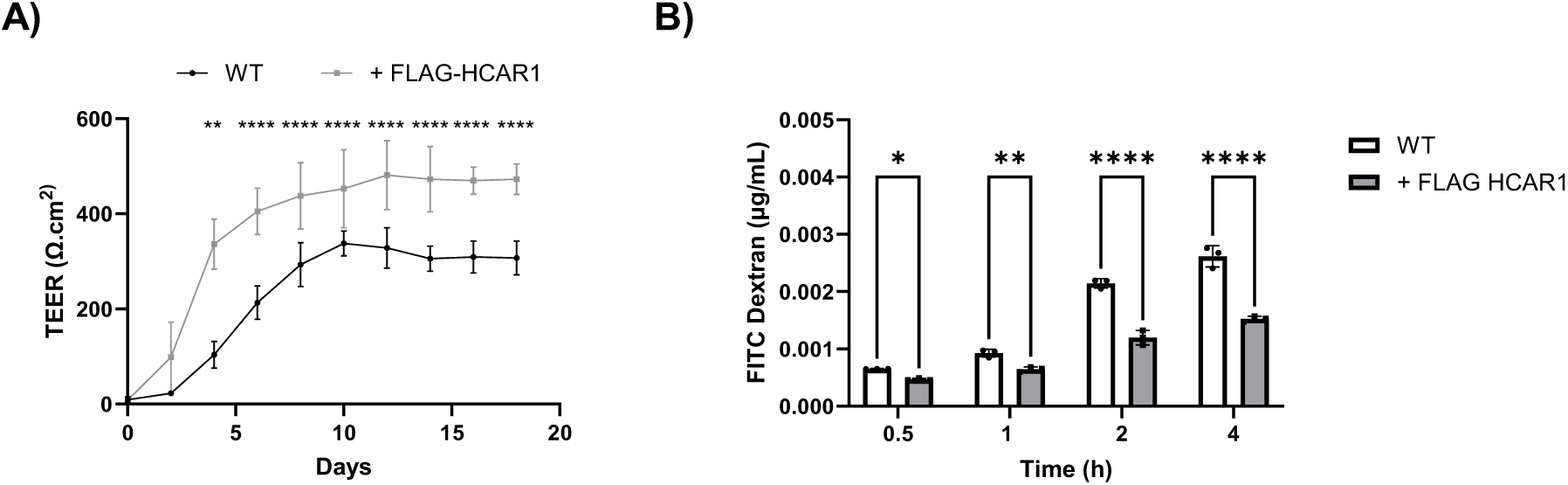
Overexpressing HCAR1 promotes cell barrier integrity and decreases cell permeability. A) TEER readings from either differentiating Caco-2 cells (black) or Caco-2 FLAG-HCAR1 (grey) cells. Caco-2 cells were grown on a 24-well Transwell membranes for 18-days with media changes every two days. TEER readings were corrected for surface area and a blank well. Data presented as mean ± SEM, N=4-6 with each Transwell condition ran in duplicates. Two-way ANOVA followed by Dunnett’s multiple comparison post-hoc test. **P<0.005, ****P<0.0001. B) Cell permeability measured using 1 mg/mL of FITC Dextran added apically and sampling across 4 h in WT Caco-2 cells (white) and Caco-2 FLAG-HCAR1 (grey) cells. Data presented as mean ± SEM, N=5. One-way ANOVA with Sidak’s multiple comparison post-hoc test. ****P<0.0001.

### Apical addition of lactate increases intestinal barrier function through a Gαi-dependent mechanism

Following the observation that increased expression of HCAR1 increased TEER readings through differentiation, WT Caco-2 cells or Caco-2 FLAG-HCAR1 cells were stimulated apically, or basally with 10 mM of either L- or D-lactate to assess the impact on intestinal barrier function. Apical addition of L- or D-lactate increased TEER readings within 10 minutes, with maximal increase in percentage change from basal occurring at 30-40 minutes. The increased TEER readings remained elevated for the duration of the experiment (Figure 5A). Greater ligand-induced differences were observed in Caco-2 FLAG-HCAR1 cells (Figure 5G). Area under the curve analysis showed that apical L-lactate addition significantly increased TEER readings compared to D-lactate, in both WT Caco-2 and Caco-2 FLAG-HCAR1 cells (Figure 5B, H). Basal addition of L-lactate did not elicit changes in TEER readings in either cell line, however, the basal addition of D-lactate resulted in a small reduction in TEER readings compared to control (Figure 5D and J). To assess if the ligand-induced alterations to cell tightness corresponded to changes in cell permeability, FITC Dextran (1 mg/mL) was added to the apical chamber and measured from the basal chamber, as a direct quantification of cellular flux. Apical addition of L- or D-lactate reduced FITC Dextran levels compared to control, suggesting that WT Caco-2 cells had reduced permeability as a result of lactate stimulation (Figure 5C). In accordance, there were greater ligand-induced differences in cell permeability observed in Caco-2 FLAG-HCAR1 cells (Figure 5I). Basal addition of L- or D-lactate did not alter levels of FITC Dextran in WT Caco-2 cells, however, basal addition of D-lactate increased cell permeability in the Caco-2 FLAG-HCAR1 cell line (Figure 5F and L).

**Figure 5.**
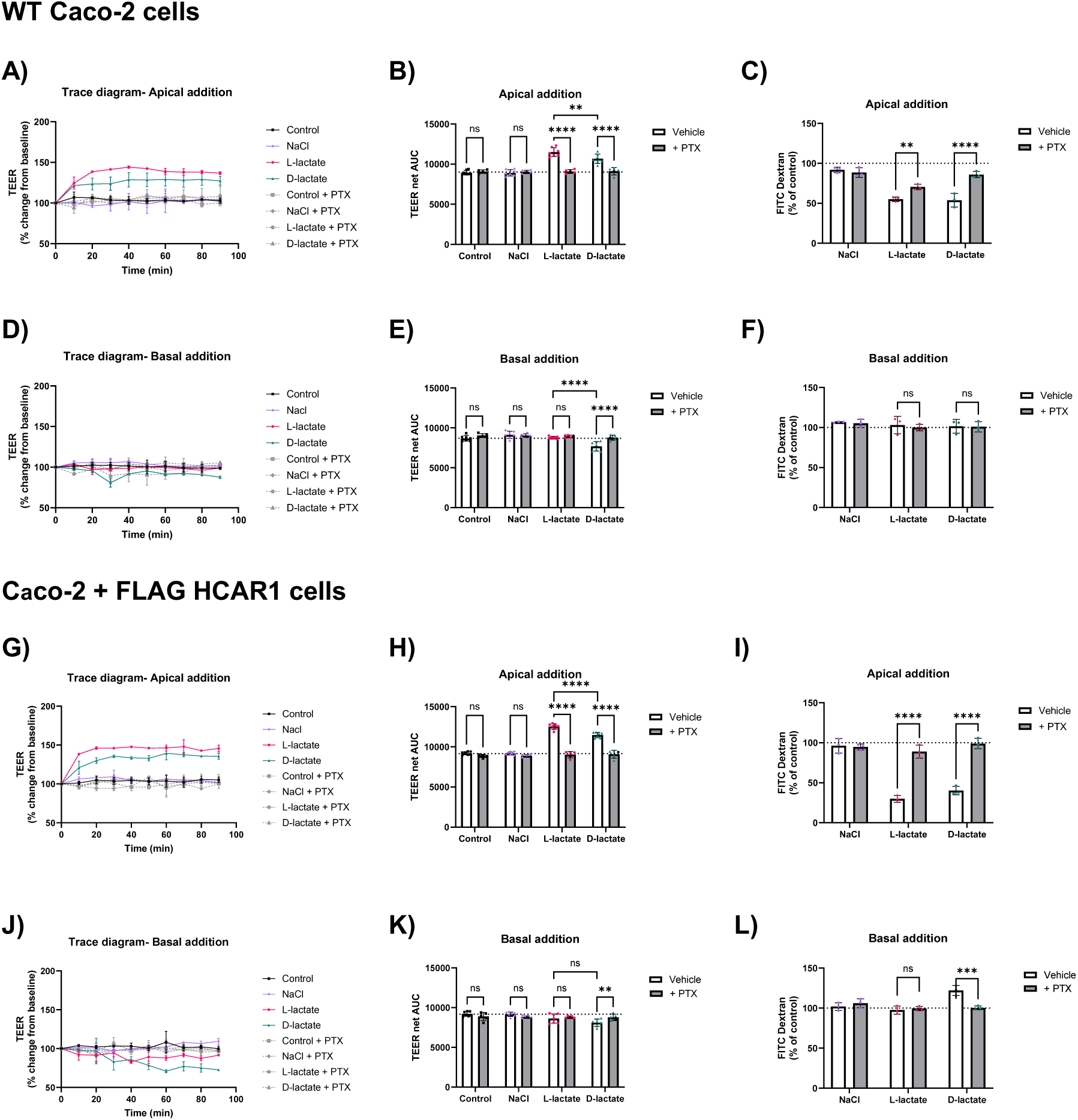
Apically added lactate enhances cell barrier integrity and reduces cell permeability. Representative traces of changes in TEER readings in response to A) apical, or D) basal addition of 10 mM lactate with or without PTX pre-treatment in differentiated WT Caco-2 cells and corresponding net AUC plotted in B) and E). Cell permeability was measured using 1 mg/mL of FITC Dextran added apically, and fluorescence intensity was measured from the basal compartment 90 minutes post C) apical or F) basal lactate stimulation. Representative traces of changes in TEER readings in response to G) apical, or J) basal addition of 10 mM lactate with or without PTX pre-treatment in differentiated Caco-2 FLAG-HCAR1 cells and corresponding net AUC plotted in H) and K). Cell permeability was measured using 1 mg/mL of FITC Dextran added apically, and fluorescence intensity was measured from the basal compartment 90 minutes post I) apical or L) basal lactate stimulation. Data presented as mean ± SEM, TEER readings were taken in triplicates and blank resistance accounted for, N=5-7. FITC Dextran measurements were taken in triplicates, N=3. Two-way ANOVA with Tukey’s post-hoc test, ****P<0.0001, ***P<0.0005, **P<0.005, ns; not significant.

Both WT Caco-2 cells and Caco-2 FLAG-HCAR1 cells were pre-treated with PTX for 16 h prior to apical or basal lactate addition. PTX attenuated the increase in apically added lactate-induced TEER readings (Figure 5B, E, H, and K) and inhibited the lactate-dependent reduction in FITC Dextran flux (Figure 5C, F, I, and L), indicating that these lactate-induced changes in epithelial cell permeability are Gαi-dependent. Interestingly, the small basal D-lactate induced reduction in TEER readings and cell permeability was also reversed by PTX (Figure 5J-L).

To determine if these lactate-dependent changes in TEER readings and cell permeability resulted in alterations in the spatial localisation of TJ proteins, the distribution and intensity profiles of ZO-1 and Claudin-1 were assessed via confocal microscopy following apical or basal lactate stimulation. Apical, but not basal, addition of either L- or D-lactate (10 mM) resulted in small punctate clusters of Claudin-1, indicated by red arrows (Figure 6A). Quantification of the fluorescence intensity of these clusters (Figure 6B) revealed that apical, but not basal, stimulation with L- or D-lactate increased Claudin-1 intensity, whereas ZO-1 remained unchanged. These findings suggest a lactate-induced rearrangement of Claudin-1 which may underlie the observed increases in TEER.

**Figure 6.**
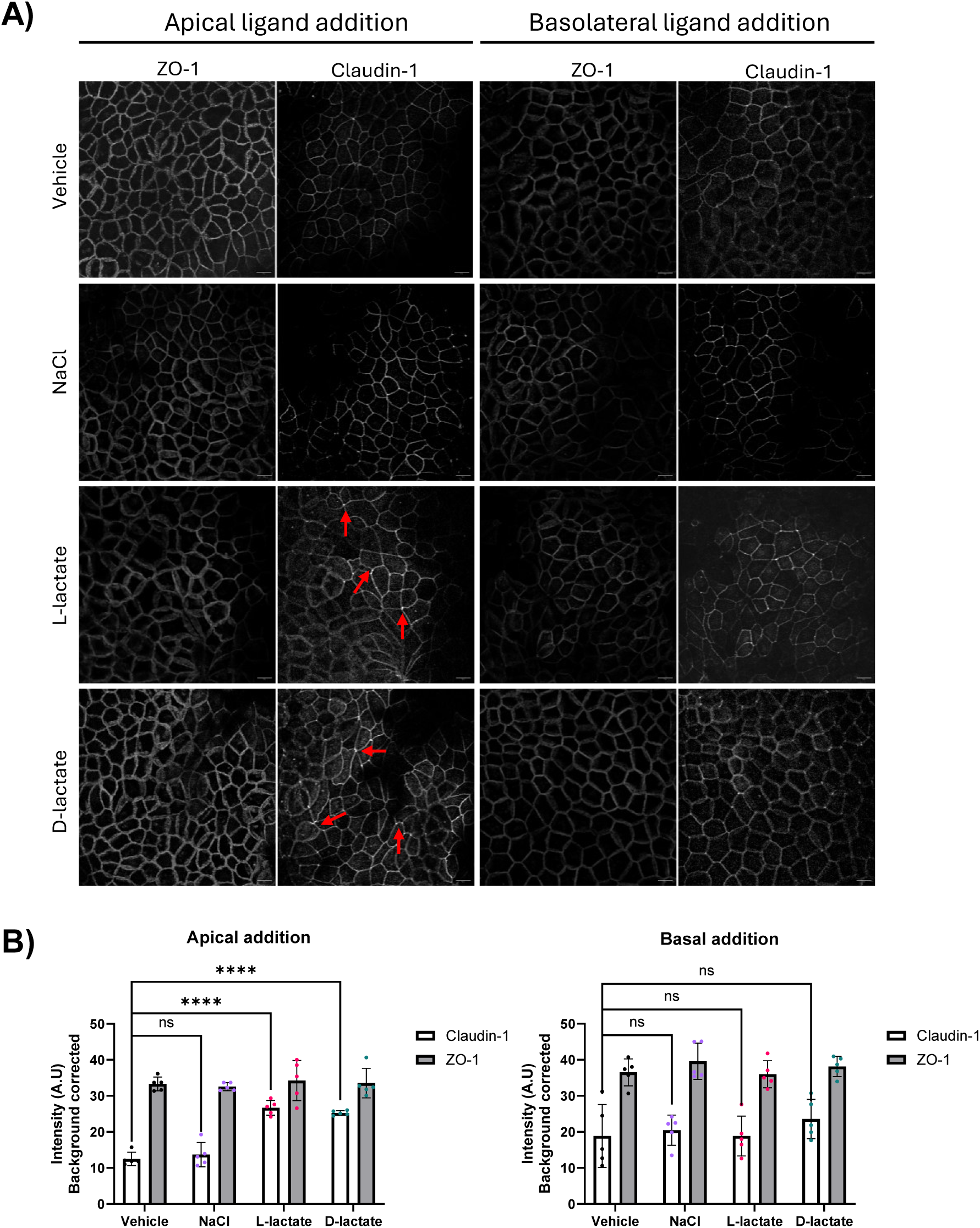
Expression profiles of ZO-1 and Claudin-1 following apical and basal lactate stimulation in differentiated Caco-2 cells. Differentiated Caco-2 cells were stimulated L- or D-lactate, NaCl (10 mM) or control, apically or basally for 90 minutes and fixed. Cells were then permeabilised and stained for ZO-1 or Claudin-1. Full Z-stacks of cells taken at same gain and intensity. Red arrows point towards protein clustering. Scale bar 10 µM. B) Fluorescence intensity plots of Claudin and ZO-1 were quantified using the multiple line intensity profile plot tool in ImageJ, corrected for background and normalised to the circumference of the cell. Data presented as mean ± SEM, N=5 where >10 individual cells were quantified for each biological repeat. One-way ANOVA, where ****P<0.0001, ns; not significant.

### Apical addition of lactate protects intestinal epithelial barrier integrity from LPS-induced damage

We next investigated if the lactate-mediated enhanced barrier function could protect against LPS-induced damage. LPS is known to be elevated in patients with inflammatory bowel disease and is commonly used to disrupt tight junctions (Guo et al., 2013). Additionally, the gut microbiota is a major source of LPS production and so closely mimics the native environment to which intestinal epithelial cells are exposed to. Caco-2 cells exposed to 1 µg/mL of LPS (*E. coli. O111:B4*) apically for 24 h significantly decreased TEER readings and increased cell permeability without inducing cell death (Supplemental Figure 3), therefore 1 µg/mL of LPS was used to disrupt tight junctions in further experiments. To assess whether L- or D-lactate could prevent LPS-induced damage, differentiated Caco-2 cells were stimulated apically or basally with L- or D-lactate for 90 minutes, followed by 1 µg/mL of LPS for 24 h (Figure 7A). Apically added lactate increased TEER over 90 minutes, however, did not reduce LPS-induced reductions in TEER readings or increases in cell permeability. Likewise, lactate added to the basolateral chamber did not alter LPS-induced damage to barrier function (Figure 7B-D). To evaluate whether L- or D-lactate could reduce LPS-induced damage, Caco-2 cells were first exposed to LPS for 24 h, followed by 90 minutes of L- or D-lactate added apically or basally. Interestingly, the addition of apical lactate after LPS exposure (Figure 7E) resulted in increased TEER readings compared to control-treated cells (Figure 7F). Coinciding with lactate-induced increased TEER readings post LPS treatment, L- or D-lactate added apically also reduced the paracellular flux of FITC Dextran post LPS stimulation (Figure 7G), indicating that cell permeability had reduced as a result of lactate addition. Consistent with our prior data (Figure 5), basolateral addition of either L- or D-lactate post LPS stimulation did not modify LPS-induced changes in the tightness of the tight junctions or cell permeability (Figure 7F, H).

**Figure 7.**
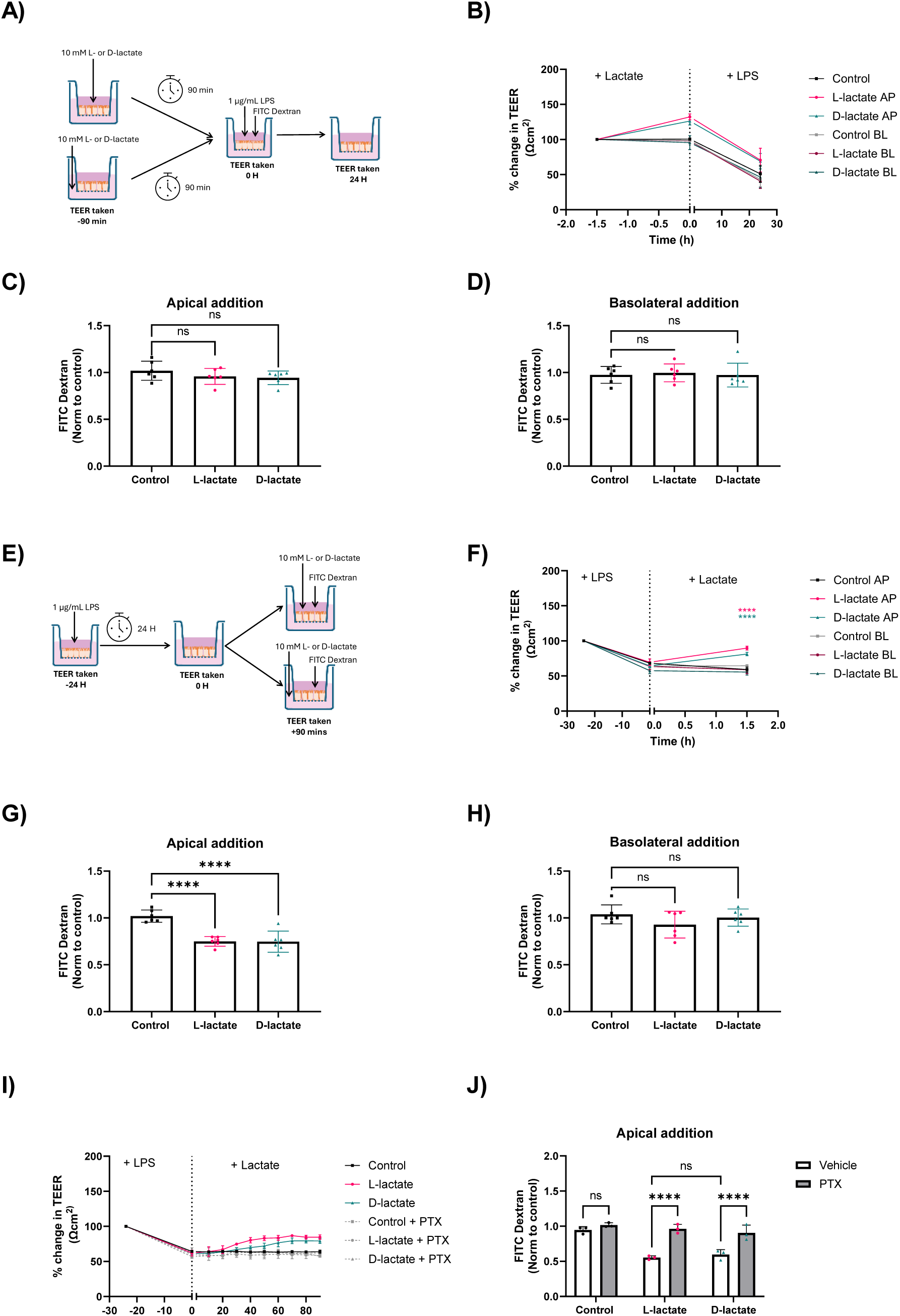
Apically added lactate reduces LPS-induced damage on barrier function in a Gαi/o-dependent manner. A) Schematic of differentiated WT Caco-2 cells pre-treated apically (AP) or basally (BL) with 10 mM lactate before LPS exposure (1 µg/mL, 24 h). Changes in B) TEER readings, and C and D) cell permeability was measured using a were measured using FITC-Dextran Flux assay. E) Schematic of differentiated WT Caco-2 cells pre-treated with LPS (1 µg/mL, 24 h) and then stimulated apically or basally with 10 mM lactate. Changes in F) TEER readings, and G and H) cell permeability was measured using a FITC-Dextran Flux assay. Data presented as mean ± SEM, TEER readings were taken in triplicates and blank resistance accounted for, N=4-6. FITC Dextran measurements were taken in triplicates, N=3. One-way ANOVA with Tukey’s post-hoc test, ****P<0.0001, ***P<0.0005, ns; not significant. I) Differentiated WT Caco-2 cells were pre-treated with LPS (1 µg/mL, 24 h), with or without PTX (200 ng/mL, 16 h) and then stimulated apically with 10 mM L- or D-lactate. TEER readings were taken every 10 minutes post LPS treatment. J) Cell permeability was measured using a FITC-Dextran Flux assay post LPS and lactate treatment. Data presented as mean ± SEM, TEER readings were taken in triplicates and blank resistance accounted for, N=3. FITC Dextran measurements were taken in triplicates, N=3. One-way ANOVA with Tukey’s post-hoc test, ****P<0.0001, ***P<0.0005, ns; not significant.

To understand the mechanisms underlying apically added lactate-mediated enhancement on barrier function, the role of Gαi-signalling was assessed. Differentiated Caco-2 cells were exposed to LPS for 24 h with and without PTX followed by a 90-minute apical treatment of L- or D-lactate and TEER readings were taken every 10 minutes. TEER increased after 20 minutes of either L- or D-lactate addition, however, post-LPS stimulation, TEER increased after 40-60 minutes compared to control, with L-lactate elevating TEER faster than D-lactate (Figure 7I). The addition of PTX prevented lactate-dependent increases in TEER. This was also reflected in cell permeability with less FITC Dextran present basolaterally post-lactate treatment compared to both control and PTX-treated cells (Figure 7J).

## Discussion

The symbiotic relationship between microbiota-derived metabolites and gastrointestinal health plays a crucial role in understanding both physiological and pathophysiological disease processes, with intestinal barrier function being paramount to the onset and progression of dysbiosis. Lactate, produced by both microbiota fermentation and during anaerobic respiration has emerged as a key metabolite for regulating energy homeostasis, metabolic health, and signalling pathways by activating HCAR1. Despite this, the mechanism by which lactate modulates HCAR1 signalling, and in turn, influences intestinal barrier function, remains poorly understood. In this study, we propose that microbiota-derived lactate enhances intestinal barrier integrity by activating apically expressed HCAR1 to increase tightening of tight junctions and reducing cell permeability, through a Gαi-dependent signalling mechanism.

We investigated HCAR1 signalling following activation by both stereoisomers of lactate. Our data revealed that L-lactate exhibited greater potency and efficacy to HCAR1 than D-lactate, which is consistent to previously reported EC50 values of L-lactate (1-5 mM) (Alessandri et al., 2024; Cai et al., 2008; Liu et al., 2009). Lactate-producing species (LAB), *lactobacilli* and *bifidobacterial species*, in the gut microbiota can produce both L- and D-lactate isoforms. Earlier studies had dismissed D-lactate as a ligand for HCAR1 (Cai et al., 2008; Liu et al., 2009), however, we demonstrated that D-lactate is capable of eliciting Gαi signalling, consistent with in vivo studies by (Órdenes et al., 2021; Scavuzzo et al., 2020). Additionally, D-lactate was found to activate ERK signalling, with a kinetic profile comparable to L-lactate. Most prior studies on HCAR1 signalling have concentrated on L-lactate as the primary physiological signalling metabolite. Yet, growing evidence indicates that D-lactate-mediated HCAR1 signalling may also have physiological relevance, justifying further exploration.

Despite HCAR1 being expressed in intestinal epithelial cells, the physiological function of this receptor in the gut was unclear. By profiling HCAR1 localisation in polarised epithelial cells, we found that the receptor was predominantly expressed on the apical membrane, alluding to the receptor responding to luminal concentrations of lactate that can been produced by LAB species from the gut microbiota. Overexpression of HCAR1 in differentiating Caco-2 cells resulted in increased TEER readings compared to WT Caco-2 cells, which was potentially indicative of the receptors’ role in maintaining/promoting intestinal barrier function. This enhancement in barrier function may be attributed to the receptor utilising endogenous lactate produced by the cells to increase expression of tight junction proteins, as previously observed *in vivo* (Li et al., 2024). Furthermore, the apical expression of HCAR1 aligns with observed apical-basal polarity in lactate responsiveness. In accordance, only lactate added apically increased barrier function and reduced paracellular flux, with more pronounced effects observed in the Caco-2 FLAG-HCAR1 overexpression system. Although intracellular transport of basolateral lactate via monocarboxylate transporters (e.g., MCT1, which is expressed on both apical and basolateral membranes) might theoretically enable indirect HCAR1 activation, this seems unlikely in our model. MCT1 has a marked preference for L-lactate over D-lactate (Halestrap, 2013; Manosalva et al., 2022), however, as we only observed a small change in TEER following basolateral D-lactate treatment, it is suggestive that MCT-mediated transport is unlikely to account for this. Additionally, we did not see an increased expression of MCT1, SMCT1 or SMCT2 in Caco-2 FLAG-HCAR1 cells compared to WT Caco-2 cells, and therefore, differences in lactate-dependent TEER readings cannot be attributed to changes in (S)MCTs. Moreover, given that D-lactate elicits less potent Gαi signalling and partial PTX-dependence in ERK signalling, the observed response may instead reflect stereoisomer-specific differences in signal pathway activation, which warrants further investigation.

Numerous microbial-derived metabolites have been reported to enhance intestinal barrier function, in Caco-2 cell models, including short-chain fatty acids, acetate (Hsieh et al., 2015) and butyrate (Peng et al., 2009), tryptophan-based metabolites (Natividad et al., 2018) and cystine (Hasegawa et al., 2021). The use of the Caco-2 cell model highlights that these metabolites can regulate barrier integrity independently of immune cell activation. LAB species such as Lactobacillus and Bifidobacterium have also been reported to increase TEER readings and promote ZO-1 and Occludin expression (Swantek et al., 1999), thus increasing intestinal barrier integrity. Furthermore, lactate’s ability to enhance epithelial barrier function is not limited to the intestine Delgado-Diaz et al., and Schwecht et al., with reports that lactic acid enhances vaginal epithelial barrier integrity (Delgado-Diaz et al., 2022; Schwecht et al., 2023).

Here, we further explored whether LPS-induced damage to barrier function could be modulated by pre- or post-addition of lactate, and whether these effects were dependent on HCAR1. LPS is known to compromise barrier integrity through activation of NF-κB and stimulate the production of pro-inflammatory cytokines (Laukoetter et al., 2008). Given prior reports that lactate suppresses LPS-induced NF-κB activation in macrophages (Hoque et al., 2014) and intestinal epithelial cells (Li et al., 2024), it is plausible that lactate, acting via HCAR1, attenuated NF-κB signalling to protect TJs from LPS-mediated damage. Consistent with this, inhibition of Gαi/o with PTX abrogated lactate-dependent increases in TEER following LPS stimulation, underscoring the importance of the lactate-HCAR1-NF-κB signalling axis. Although a pre-treatment of apically added lactate increased TEER, it did not prevent LPS-induced damage to the tight junctions, indicating that increasing levels of microbiota-derived lactate would not prevent intestinal barrier damage from occurring. Instead, post-induction of damage may be rectified with acute increases in microbiota-derived lactate. These findings are in accordance with Iraporda et al., and Li et al., who demonstrated that lactate could offer protection against experimental-induced colitis in murine models (Iraporda et al., 2016; Li et al., 2024).

Overall, this work has alluded towards lactate-dependent mechanisms in enhancing intestinal barrier function by promoting the tightening of TJs and reducing cell permeability. Although there were no differences in the way the stereoisomers acted on intestinal barrier function, there was a preferential signalling sidedness to the response, with apical (luminal) addition of lactate being favourable. This coincides with HCAR1 being predominantly expressed at the apical membrane. The protective effect of lactate on the intestinal barrier function was still observed following LPS-induced damage, and consequently, could hold therapeutic potential for people with inflammatory bowel diseases.

Limitations: While this study utilises the human Caco-2 cell model to investigate the role of lactate in intestinal epithelial barrier integrity, we acknowledge several limitations of this system. These include the overestimation of TEER values compared to in vivo models (Ebert et al., 2024), the absence of continuous fluid flow and fluid shear stress (Delon et al., 2019), and the lack of mucus-secreting cells (Cheng et al., 2023), which alters metabolite absorption. Recent advances in complex in vitro systems offer more physiologically relevant alternatives, including microfluidic platforms (Tan et al., 2018), organoids and mini-colons (Mitrofanova et al., 2024), and gut-on-a-chip models (Gleeson et al., 2024), which could be employed to better characterise the effects of lactate on barrier function. Furthermore, the lack of commercially available selective receptor antagonists limits the ability to assess the direct role of acute HCAR1 activation. Although CRISPR-based knockout models could be developed to address this, such genetic modifications carry the risk of inadvertently altering other cellular functions.

## Methods

### Plasmid construct

A stop codon (TGA) at position 809 was introduced to FLAG-HCAR1-Tango plasmid (Addgene #66395) to prevent the transcription of Tango. Mutagenic primers were designed using software on the Agilent website and PCR conducted.

### Cell culture

Caco-2 cells (C2BBe1 clonal) were obtained from the American Type culture collection and routinely maintained as a monolayer in T-75 flasks at 37 °C in a 5 % CO_2_ humidified incubator. Cells were cultured in Dulbecco’s Modified Eagle’s Medium (DMEM) media (Gibco) containing 4.5 g/L glucose, 1% glutamine and supplemented with 10 % fetal bovine serum (Gibco) and 1 % (v/v) streptomycin/penicillin (Sigma). Cells were used between passage 78-82.

Caco-2 cells stably expressing FLAG-HCAR1 were generated by transfecting Caco-2 cells with FLAG-HCAR1 plasmid containing G418 resistance gene for 48 h using Lipofectamine 2000 reagent (Invitrogen). Cells were then subjected to G418 exposure (1 mg/mL) for two weeks. Surviving cells were isolated and sorted by FACs to form polyclonal populations. The population of Caco-2 cells highly expressing FLAG-HCAR1 were used in these results.

Caco-2 cells were grown in monolayers on either 6-well or 24-well transwell inserts, 0.4 µM pore size (Corning) to enable differentiation into polarised epithelial cells. Cells were seeded at a density of 10^5^ cells per cm^2^ on the filter membranes, and apical and basolateral chambers were bathed in media. The media was refreshed every two days for 18 days. Assays were conducted on day 18 of differentiation.

### Transepithelial Electrical Resistance (TEER) readings

Transepithelial electrical resistance (TEER) readings measure the integrity of barrier formation and are a strong indicator of differentiated Caco-2 cells. TEER readings were taken every two days for 18 days, until the reading had stabilised, using EVOM3 TEER machine. Briefly, STX2-PLUS electrode probes were sterilised with 70 % ethanol, allowed to air dry, and calibrated to complete media. The probes were placed into the Transwell chamber perpendicularly to the plate, with one electrode entering the basolateral compartment and the other electrode entering the apical compartment without touching the monolayer of cells. Resistance across the membrane insert was measured in triplicates. Readings were corrected for background resistance due to the membrane insert and collagen layer, and surface area.

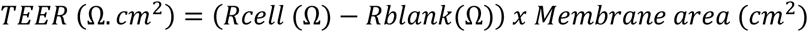

### Paracellular Flux assay

Paracellular flux was determined using 4 KDa FITC Dextran (Sigma) at a final concentration of 1 mg/mL. Following differentiation, 100 µL of FITC Dextran was added into the apical chamber and three parallel 100 μL samples were collected from the basolateral chamber of the Transwell post-treatment and added directly to a 96-well black plate. Equivalent media was added back into the basolateral chamber to prevent bulk fluid flow imbalance. The fluorescence emission at 535 nm was measured after excitation at 475 nm by using a LUMIstarOPTIMA plate reader (BMG Labtech). Concentration of basolateral FITC Dextran was determined through extrapolation of a standard curve of predetermined concentrations. Paracellular permeability coefficients (P_app_) were calculated using the principle below:

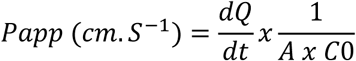

Where dQ/dt is the temporal mass flow into the basal chamber, A is the area of transwell and C_0_ is the initial concentration of product added to apical chamber.

### LPS Stimulation

Lipopolysaccharide (LPS) derived from E. coli. O111:B4 (Sigma) was reconstituted in water to a concentration of 1 mg/mL and stored in aliquots at −20 °C. Upon use, 1 mg/mL LPS aliquot was dilute to 10 μg/mL and stored in a silanized glass container at 4 °C for one month. LPS was added to the apical side of differentiated Caco-2 cells at a final concentration of 1 μg/mL for 24 h.

### cAMP assay

#### cAMP accumulation assay

Caco-2 cells were seeded in triplicate format at a density of 40,000 cells/well into a 96-well plate and grown to 90 % confluency. Cells were washed with PBS and pre-treated with 0.5 mM IBMX diluted in DMEM for 5 minutes at 37 °C. Ligands were diluted to desired concentration in 0.5 mM IBMX containing DMEM supplemented with 3 µM forskolin and added to cells for 5 minutes at 37 °C. Following stimulation, cells were washed with ice-cold PBS and lysed. cAMP levels were determined using cAMP Gs Dynamic 2 kit (Revvity) and normalised to protein concentration.

#### GloSensor cAMP assay

Caco-2 cells were grown in a 6-well plate to 80 % confluency and transfected with 3000 ng of p-GloSensor-20F plasmid for 24 h using Lipofectamine 2000. 14 h post-transfection, Caco-2 cells were replated at a density of 50,000 cells/well into a 96-well white plate and allowed to adhere for 8 h. DMEM media was replaced with equilibrium buffer (88 % CO2-independent media, 10 % FBS and 2 % GloSensor) and incubated at 37 °C for 2 h. The 96-well white plate was then placed into a 37 °C pre heated PHERAstar FSX plate reader and basal luminescence conditions were determined over 10 cycles (1 cycle, 10 seconds). Forskolin at a final concentration of 3 µM (diluted in equilibrium buffer) was added directly to the wells and luminescence detected for 5 minutes. At this point, lactate (various concentrations, diluted in equilibrium buffer) was added to wells and the plate was read for a further 25 minutes. Data were normalised to the basal luminescence read and then to the individual well’s forskolin response.

### Flow cytometry

Flow cytometry was used to quantitate receptor uptake and cell surface expression. Caco-2 FLAG HCAR1 cells were split into two wells and grown to 70% confluency. One set of cells were then incubated with M1 anti-FLAG-antibody (1:500, Sigma) diluted in serum free DMEM and the second set of cells were incubated with serum free DMEM for 20 minutes at 37 °C. If cells were being treated with ligand, this was added after the M1 treatment. Cells were then gently washed using ice-cold PBS/ Ca2+, harvested in FACs buffer (ice-cold PBS / 2 % FBS) and centrifuged at 1000 rpm at 4 °C for 10 minutes. The supernatant was removed, and the pellet was resuspended in FACs buffer containing AlexaFluor 488 anti-mouse secondary antibody (1:1000) and incubated for 1 h on ice. A subset of cells were resuspended in FACs buffer only to allow for normalisation to basal cell fluorescence. Following secondary antibody labelling, cells were centrifuged at 1000 rpm at 4 °C for 10 minutes and resuspended with 1 mL of FACs buffer. This was repeated 3 more times with the final resuspension volume of 300 µL of FACs buffer. The suspended cells were transferred into round bottomed polypropylene FACs tubes and analysed on a FACs Calibur Flow cytometer.

### qPCR

Total RNA was extracted from Caco-2 cells using TRIzol reagent (Invitrogen), purified and treated with an RNAse inhibitor (Thermofisher) and DNase I treatment (Life Technologies). 1 µg of RNA was converted to cDNA according to manufacturer’s guidelines using SuperScript IV Reverse Transcriptase kit (Life Technologies). Briefly, cDNA, forward and reverse primers, 10 µL of 1X SYBR-Green PCR Mastermix and nuclease-free water to a total volume of 20 µL were added in a white-bottom 96 well plate. Each reaction was run in triplicates along with a negative control replacing cDNA with nuclease-free water. Reaction were performed using StepOnePlus Applied Biosystems machine. Briefly, the sample was denatured using cycles at 50 °C for 10 mins and 95 °C for 10 minutes, followed by 50 cycles of annealing and extension: 95 °C for 15 seconds, 60 °C for 42 seconds, and then dissociation at 95 °C for 15 seconds, 60 °C for 60 seconds and 95 °C for 15 seconds. The expression level of each gene was determined using the ΔCT method and normalised to the GAPDH housekeeping gene. For comparison of time points or conditions 2^-ΔΔCT^ method were used. Primers were generated through NIH-BLAST tool and are listed in Supplementary table 1. qPCR primer design was performed using primer-BLAST. The amplicon for each primer set that was generated was checked for <60 % GC content, and the predicted derivative plot and predicted melt curves using uMELT Quartz software. Finally, the primer pair were checked for specificity against the target sequence to ensure unspecific targets were not going to be amplified.

### Western blotting

Proteins were extracted from cell lysates using ice-cold RIPA lysis buffer (50 mM Tris pH 7.4, 1 % Triton X-100, 140 mM NaCl, 5 mM EDTA, 1 mM NaF, 1 mM PMSF, 1 mM Na_3_VO_4_ and a protease inhibitor tablet (Roche)). Protein concentrations were determined using Pierce Coomassie (Bradford) Protein Assay kit and extrapolated from a BSA standard curve and diluted in RIPA buffer to standardise concentration. Laemmli sample buffer with 5 % 2-β-mercaptoethanol was added to diluted sample and heat shocked at 95 °C for 5 minutes. Samples were then separated with 12 % SDS-polyacrylamide gel electrophoresis and transferred to nitrocellulose membrane. The membrane was blocked and probed for Occludin, Claudin-1, p-p65, p65 and α-tubulin with corresponding primary antibodies and horseradish peroxidase-conjugated secondary antibodies. Membranes were imaged using Immobilon Forte Western HRP reagent (Millipore) and Chemi-imager. Band intensities were analysed using image J.

### Confocal microscopy

Undifferentiated Caco-2 cells grown on glass coverslips, were and fed anti-FLAG antibody (1:500) live for 20 minutes at 37 °C followed by ligand stimulation. Cells were washed 3x with ice-cold PBS/ Ca^2+^ and if required, stripped using 0.04 M EDTA, and fixed with 4% PFA (w/v) for 20 minutes at RT. Fixed cells were blocked in PBS/Ca^2+^ with 2 % FBS for 1 h at RT. Cells were washed a further three times with PBS before incubation with Alexa Fluro-conjugated secondary antibody (1:1000, Thermofisher) for 1 h at RT. Coverslips were carefully mounted onto slides using Fluormount-G containing DAPI, and imaged. Differentiated Caco-2 cells grown on filter membrane transwell were treated as described above. If further primary antibodies were required, cells were permeabilised for 15 minutes with 0.2 % Triton-X in PBS at RT, followed by overnight incubation of primary antibody diluted to desired concentration in blocking buffer at 4 °C. For mounting, membranes were excised out of the Transwell using scalpels and tweezers, before being placed apical side up-right onto a glass slide. A coverslip with Fluoromount-G with DAPI was then placed on top of the apical side of the membrane and sealed.

Images were acquired with Leica Stellaris 8 microscope using the 63x oil-immersion objective at excitation wavelengths of 455, 488 and 647 nm. Gain and offset were corrected at the beginning of an experiment and were kept constant between conditions to allow for comparison. Images acquired in lightning mode had 0.5 AU pinhole, a line average of 4, and an optical section of approximately 0.4 μM. Full Z-stacks of differentiated Caco-2 cells were acquired.

Analysis was conducted using the multiple line intensity profile plot tool in Image J. Fluorescence intensity was normalised to the diameter or circumference of the cell being analysed and corrected for background.

### Quantification and statistical analysis

All statistical analysis was performed in GraphPad Prism version 10, with specific tests noted in each figure legend. Data are presented as either raw values or as a fold-change relative to the most informative comparator (either basal or control). One-way ANOVA with Tukey’s or Dunnett’s post-hoc test was used for multiple comparisons for groups of 3 or more. Two-way ANOVA was used to compare two groups under multiple conditions. All experiments were repeated at least three times with actual N numbers reported in figure legends. In all cases, P < 0.05 were considered significant.

## Acknowledgements

We would like to acknowledge Dr Stephen Rothery at the Facility for Imaging of Light Microscopy at Imperial College London for his technical support for microscopy-based experiments, and Dr Rachael Barry for providing the Caco-2 cell line. Graphical schematics were produced using Biorender. This project was funded by the Medical Research Council and Sosei Heptares MultiSci iCASE (MR/R015732/1) and Genesis Research Trust.

## Author Contributions

A.J.M, G.S.F and A.C.H conceived and designed the experiments. A.J.M performed the majority of laboratory experiments, analysis and figure preparation and with P.A conducted differentiation of cells. A.J.M and A.C.H wrote the manuscript with inputs and approval of the final manuscript from all authors.

## References

Alessandri, M., Osorio-Forero, A., Lüthi, A., & Chatton, J.-Y. (2024). The lactate receptor HCAR1: A key modulator of epileptic seizure activity. IScience, 27(5), 109679. 10.1016/j.isci.2024.109679

Brooks, G. A. (2021). Role of the Heart in Lactate Shuttling. Frontiers in Nutrition, Volume 8*-* 2021. https://www.frontiersin.org/journals/nutrition/articles/10.3389/fnut.2021.663560

Cai, T.-Q., Ren, N., Jin, L., Cheng, K., Kash, S., Chen, R., Wright, S. D., Taggart, A. K. P., & Waters, M. Gerard. (2008). Role of GPR81 in lactate-mediated reduction of adipose lipolysis. Biochemical and Biophysical Research Communications, 377(3), 987–991. 10.1016/j.bbrc.2008.10.088

Cheng, Y., Watanabe, C., Ando, Y., Kitaoka, S., Egawa, Y., Takashima, T., Matsumoto, A., & Murakami, M. (2023). Caco-2 Cell Sheet Partially Laminated with HT29-MTX Cells as a Novel In Vitro Model of Gut Epithelium Drug Permeability. Pharmaceutics, 15(9), 2338. 10.3390/pharmaceutics15092338

Delgado-Diaz, D. J., Jesaveluk, B., Hayward, J. A., Tyssen, D., Alisoltani, A., Potgieter, M., Bell, L., Ross, E., Iranzadeh, A., Allali, I., Dabee, S., Barnabas, S., Gamieldien, H., Blackburn, J. M., Mulder, N., Smith, S. B., Edwards, V. L., Burgener, A. D., Bekker, L.-G., … Tachedjian, G. (2022). Lactic acid from vaginal microbiota enhances cervicovaginal epithelial barrier integrity by promoting tight junction protein expression. Microbiome, 10(1), 141. 10.1186/s40168-022-01337-5

Delon, L. C., Guo, Z., Oszmiana, A., Chien, C.-C., Gibson, R., Prestidge, C., & Thierry, B. (2019). A systematic investigation of the effect of the fluid shear stress on Caco-2 cells towards the optimization of epithelial organ-on-chip models. Biomaterials, 225, 119521. 10.1016/j.biomaterials.2019.119521

Ebert, A., Dahley, C., & Goss, K.-U. (2024). Pitfalls in evaluating permeability experiments with Caco-2/MDCK cell monolayers. European Journal of Pharmaceutical Sciences, 194, 106699. 10.1016/j.ejps.2024.106699

Feingold, K. R., Moser, A., Shigenaga, J. K., & Grunfeld, C. (2011). Inflammation inhibits GPR81 expression in adipose tissue. Inflammation Research, 60(10), 991–995. 10.1007/s00011-011-0361-2

Feng, J., Yang, H., Zhang, Y., Wei, H., Zhu, Z., Zhu, B., Yang, M., Cao, W., Wang, L., & Wu, Z. (2017). Tumor cell-derived lactate induces TAZ-dependent upregulation of PD-L1 through GPR81 in human lung cancer cells. Oncogene, 36(42), 5829–5839. 10.1038/onc.2017.188

Ghosh, S., Whitley, C. S., Haribabu, B., & Jala, V. R. (2021). Regulation of Intestinal Barrier Function by Microbial Metabolites. Cellular and Molecular Gastroenterology and Hepatology, 11(5), 1463–1482. 10.1016/j.jcmgh.2021.02.007

Gleeson, J. P., Zhang, S. Y., Subelzu, N., Ling, J., Nissley, B., Ong, W., Nofsinger, R., & Kesisoglou, F. (2024). Head-to-Head Comparison of Caco-2 Transwell and Gut-on-a-Chip Models for Assessing Oral Peptide Formulations. Molecular Pharmaceutics, 21(8), 3880–3888. 10.1021/acs.molpharmaceut.4c00210

Guo, S., Al-Sadi, R., Said, H. M., & Ma, T. Y. (2013). Lipopolysaccharide Causes an Increase in Intestinal Tight Junction Permeability in Vitro and in Vivo by Inducing Enterocyte Membrane Expression and Localization of TLR-4 and CD14. The American Journal of Pathology, 182(2), 375–387. 10.1016/j.ajpath.2012.10.014

Halestrap, A. P. (2013). The SLC16 gene family – Structure, role and regulation in health and disease. Molecular Aspects of Medicine, 34(2–3), 337–349. 10.1016/j.mam.2012.05.003

Hasegawa, T., Mizugaki, A., Inoue, Y., Kato, H., & Murakami, H. (2021). Cystine reduces tight junction permeability and intestinal inflammation induced by oxidative stress in Caco-2 cells. Amino Acids, 53(7), 1021–1032. 10.1007/s00726-021-03001-y

Hoque, R., Farooq, A., Ghani, A., Gorelick, F., & Mehal, W. Z. (2014). Lactate Reduces Liver and Pancreatic Injury in Toll-Like Receptor– and Inflammasome-Mediated Inflammation via GPR81-Mediated Suppression of Innate Immunity. Gastroenterology, 146(7), 1763–1774. 10.1053/j.gastro.2014.03.014

Horowitz, A., Chanez-Paredes, S. D., Haest, X., & Turner, J. R. (2023). Paracellular permeability and tight junction regulation in gut health and disease. Nature Reviews Gastroenterology & Hepatology, 20(7), 417–432. 10.1038/s41575-023-00766-3

Hsieh, C.-Y., Osaka, T., Moriyama, E., Date, Y., Kikuchi, J., & Tsuneda, S. (2015). Strengthening of the intestinal epithelial tight junction by *Bifidobacterium bifidum*. Physiological Reports, 3(3), e12327. 10.14814/phy2.12327

Iraporda, C., Romanin, D. E., Bengoa, A. A., Errea, A. J., Cayet, D., Foligné, B., Sirard, J.-C., Garrote, G. L., Abraham, A. G., & Rumbo, M. (2016). Local Treatment with Lactate Prevents Intestinal Inflammation in the TNBS-Induced Colitis Model. Frontiers in Immunology, 7. 10.3389/fimmu.2016.00651

Johannessen, L. E., Spilsberg, B., Wiik-Nielsen, C. R., Kristoffersen, A. B., Holst-Jensen, A., & Berdal, K. G. (2013). DNA-Fragments Are Transcytosed across CaCo-2 Cells by Adsorptive Endocytosis and Vesicular Mediated Transport. PLOS ONE, 8(2), e56671-. 10.1371/journal.pone.0056671

Laukoetter, M. G., Nava, P., & Nusrat, A. (2008). Role of the intestinal barrier in inflammatory bowel disease. World Journal of Gastroenterology, 14(3), 401. 10.3748/wjg.14.401

Lea, T. (2015). Caco-2 Cell Line. In The Impact of Food Bioactives on Health (pp. 103–111). Springer International Publishing. 10.1007/978-3-319-16104-4_10

Lee, Y.-S., Kim, T.-Y., Kim, Y., Lee, S.-H., Kim, S., Kang, S. W., Yang, J.-Y., Baek, I.-J., Sung, Y. H., Park, Y.-Y., Hwang, S. W., O, E., Kim, K. S., Liu, S., Kamada, N., Gao, N., & Kweon, M.-N. (2018). Microbiota-Derived Lactate Accelerates Intestinal Stem-Cell-Mediated Epithelial Development. Cell Host & Microbe, 24(6), 833–846.e6. 10.1016/j.chom.2018.11.002

Li, X., Yao, Z., Qian, J., Li, H., & Li, H. (2024). Lactate Protects Intestinal Epithelial Barrier Function from Dextran Sulfate Sodium-Induced Damage by GPR81 Signaling. Nutrients, 16(5), 582. 10.3390/nu16050582

Liu, C., Wu, J., Zhu, J., Kuei, C., Yu, J., Shelton, J., Sutton, S. W., Li, X., Yun, S. J., Mirzadegan, T., Mazur, C., Kamme, F., & Lovenberg, T. W. (2009). Lactate Inhibits Lipolysis in Fat Cells through Activation of an Orphan G-protein-coupled Receptor, GPR81. Journal of Biological Chemistry, 284(5), 2811–2822. 10.1074/jbc.M806409200

Manosalva, C., Quiroga, J., Hidalgo, A. I., Alarcón, P., Anseoleaga, N., Hidalgo, M. A., & Burgos, R. A. (2022). Role of Lactate in Inflammatory Processes: Friend or Foe. Frontiers in Immunology, 12. 10.3389/fimmu.2021.808799

Mitrofanova, O., Nikolaev, M., Xu, Q., Broguiere, N., Cubela, I., Camp, J. G., Bscheider, M., & Lutolf, M. P. (2024). Bioengineered human colon organoids with in vivo-like cellular complexity and function. Cell Stem Cell, 31(8), 1175–1186.e7. 10.1016/j.stem.2024.05.007

Natividad, J. M., Agus, A., Planchais, J., Lamas, B., Jarry, A. C., Martin, R., Michel, M.-L., Chong-Nguyen, C., Roussel, R., Straube, M., Jegou, S., McQuitty, C., Le Gall, M., da Costa, G., Lecornet, E., Michaudel, C., Modoux, M., Glodt, J., Bridonneau, C., … Sokol, H. (2018). Impaired Aryl Hydrocarbon Receptor Ligand Production by the Gut Microbiota Is a Key Factor in Metabolic Syndrome. Cell Metabolism, 28(5), 737–749.e4. 10.1016/j.cmet.2018.07.001

Órdenes, P., Villar, P. S., Tarifeño-Saldivia, E., Salgado, M., Elizondo-Vega, R., Araneda, R. C., & García-Robles, M. A. (2021). Lactate activates hypothalamic POMC neurons by intercellular signaling. Scientific Reports, 11(1), 21644. 10.1038/s41598-021-00947-7

Peng, L., Li, Z.-R., Green, R. S., Holzmanr, I. R., & Lin, J. (2009). Butyrate Enhances the Intestinal Barrier by Facilitating Tight Junction Assembly via Activation of AMP-Activated Protein Kinase in Caco-2 Cell Monolayers. The Journal of Nutrition, 139(9), 1619–1625. 10.3945/jn.109.104638

Pohanka, M. (2020). D-Lactic Acid as a Metabolite: Toxicology, Diagnosis, and Detection. BioMed Research International, 2020(1), 3419034. 10.1155/2020/3419034

Ranganathan, P., Shanmugam, A., Swafford, D., Suryawanshi, A., Bhattacharjee, P., Hussein, M. S., Koni, P. A., Prasad, P. D., Kurago, Z. B., Thangaraju, M., Ganapathy, V., & Manicassamy, S. (2018). GPR81, a Cell-Surface Receptor for Lactate, Regulates Intestinal Homeostasis and Protects Mice from Experimental Colitis. The Journal of Immunology, 200(5), 1781–1789. 10.4049/jimmunol.1700604

Rao, S. S. C., Yu, S., Tetangco, E. P., & Yan, Y. (2018). Probiotics can Cause D-Lactic Acidosis and Brain Fogginess: Reply to Quigley et al. Clinical and Translational Gastroenterology, 9(11), e207. 10.1038/s41424-018-0077-5

Scavuzzo, C. J., Rakotovao, I., & Dickson, C. T. (2020). Differential effects of L- and D-lactate on memory encoding and consolidation: Potential role of HCAR1 signaling. Neurobiology of Learning and Memory, 168, 107151. 10.1016/j.nlm.2019.107151

Schwecht, I., Nazli, A., Gill, B., & Kaushic, C. (2023). Lactic acid enhances vaginal epithelial barrier integrity and ameliorates inflammatory effects of dysbiotic short chain fatty acids and HIV-1. Scientific Reports, 13(1), 20065. 10.1038/s41598-023-47172-y

Srinivasan, B., Kolli, A. R., Esch, M. B., Abaci, H. E., Shuler, M. L., & Hickman, J. J. (2015). TEER Measurement Techniques for In Vitro Barrier Model Systems. SLAS Technology, 20(2), 107–126. 10.1177/2211068214561025

Swantek, J. L., Christerson, L., & Cobb, M. H. (1999). Lipopolysaccharide-induced Tumor Necrosis Factor-α Promoter Activity Is Inhibitor of Nuclear Factor-κB Kinase-dependent. Journal of Biological Chemistry, 274(17), 11667–11671. 10.1074/jbc.274.17.11667

Tan, H.-Y., Trier, S., Rahbek, U. L., Dufva, M., Kutter, J. P., & Andresen, T. L. (2018). A multi-chamber microfluidic intestinal barrier model using Caco-2 cells for drug transport studies. PLOS ONE, 13(5), e0197101. 10.1371/journal.pone.0197101

Yang, F., Chen, G., Ma, M., Qiu, N., Zhu, L., & Li, J. (2018). Fatty acids modulate the expression levels of key proteins for cholesterol absorption in Caco-2 monolayer. Lipids in Health and Disease, 17(1), 32. 10.1186/s12944-018-0675-y

