## Supplemental Figure for "D- and L-Lactate enhance intestinal barrier function via activation of an apical HCAR1/Gαi pathway in a human colonic epithelial cell model"

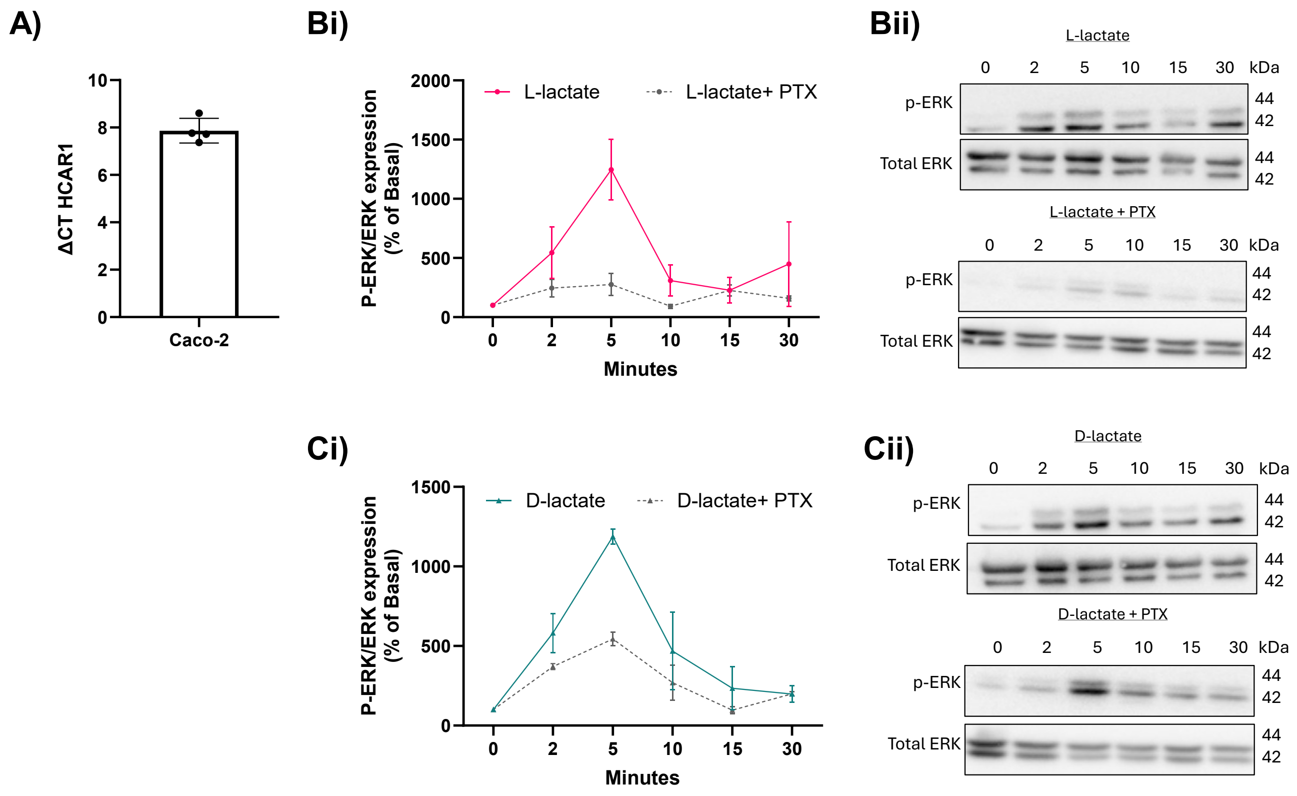


**Supplemental Figure 1: Profiling HCAR1 signalling in Caco-2 cells.** A) Confirmation of HCAR1 expression in Caco-2 cells. Caco-2 cells were grown to confluency and harvested. qPCR analysis was conducted to assess HCAR1 mRNA expression. Levels were normalised to the housekeeping gene GAPDH (ΔCT). Data presented as mean ± SEM, N=4. Phospho-ERK activity in Caco-2 cells with or without PTX pre-treatment in response to B) L- or C) D-lactate stimulation (10 mM, 0, 2, 5,10, 15 or 30 minutes). Data normalised to total-ERK and presented as percentage change of ERK activation from basal levels. Grey dotted line is response in cells pre-treated with PTX for 16 h. Data presented as mean ± SEM. N=5 for lactate stimulation, N=3 for PTX and lactate stimulation. Data presented as mean ± SEM, N=3. One-way ANOVA, with Tukey's post-hoc test, ****P<0.0001,***P<0.0005, **P<0.005.

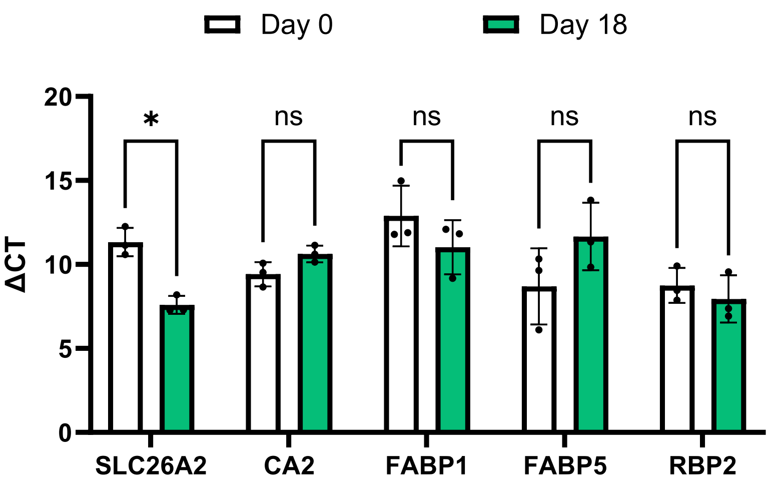


**Supplemental Figure 2: Expression of enterocyte markers in differentiating Caco-2 cells.** Undifferentiated (white) or differentiated (green) Caco-2 cells were grown to confluency and harvested. qPCR analysis was conducted to assess expression of SLC26A2, CA2, FABP1, FABP5 and RBP2 mRNA. Levels normalised to the house keeping gene GAPDH. Data presented as mean ± SEM, N=3. Unpaired two tailed Student's T-test, *P<0.05, ns; not significant.

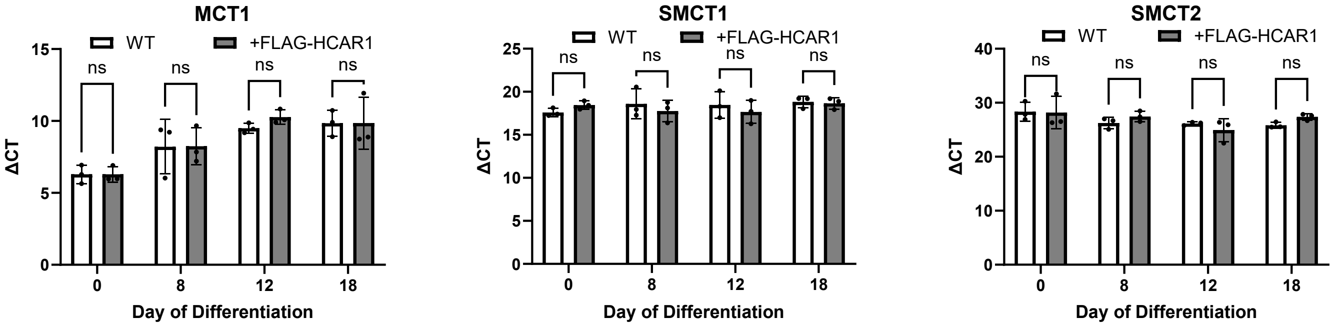


**Supplemental Figure 3: Expression of monocarboxylic acid transporters in differentiating Caco-2 cells.** Differentiating WT Caco-2 cells (white) and Caco-2 FLAG-HCAR1 (grey) cells were harvested on day 0, 8, 12 and 18 of differentiation and qPCR analysis was conducted to assess expression of MCT1, SMCT1 and SMCT2 mRNA. Levels normalised to the house keeping gene GAPDH. Data presented as mean ± SEM, N=3. Unpaired two tailed Student's T-test. ns; not significant.

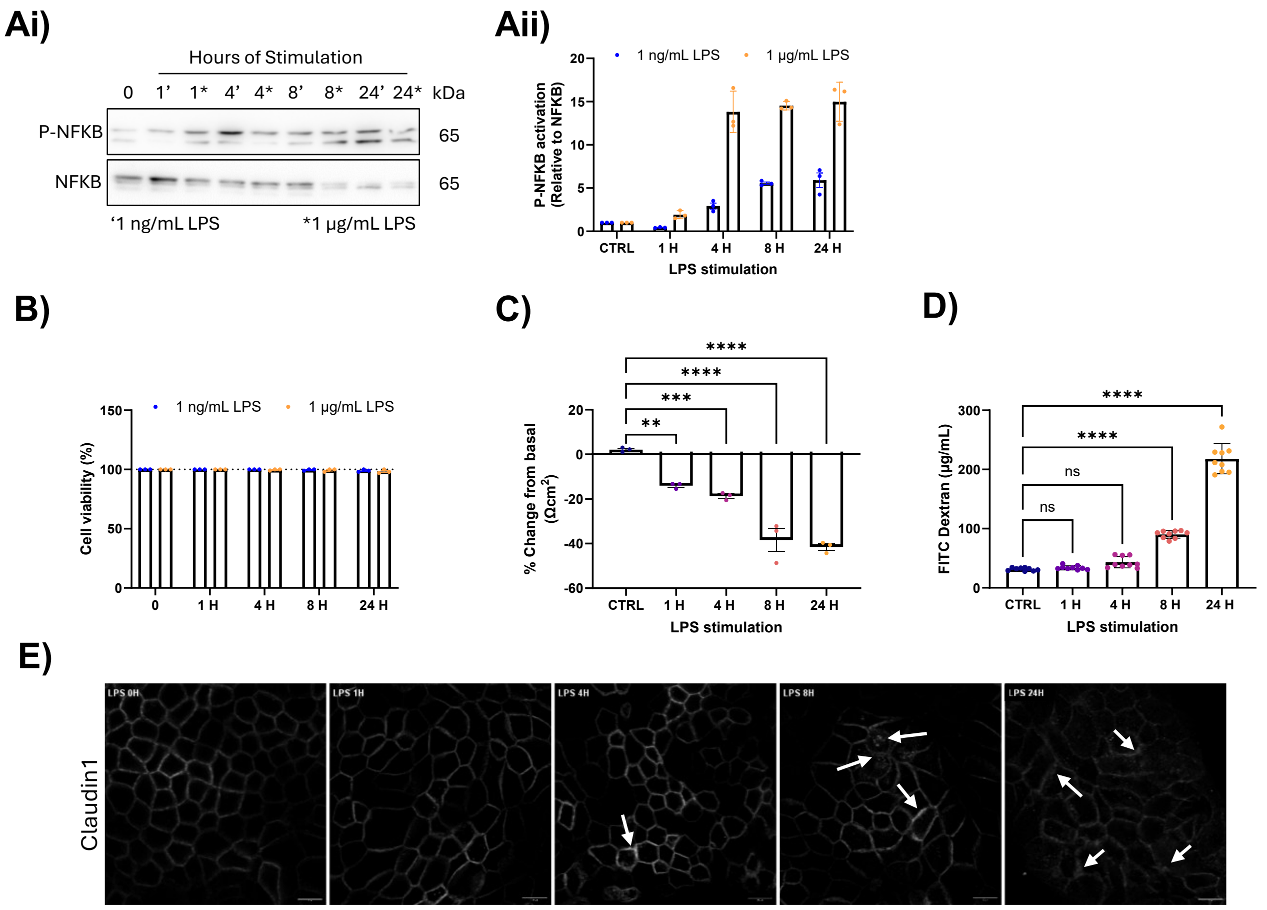


**Supplemental Figure 3: The effect of LPS stimulation on Caco-2 cells**. Undifferentiated Caco-2 cells were stimulated with 1 ng/mL or 1 µg/mL of LPS for 0, 1, 4, 8 or 24 h. Levels of p65 activity were measured via western blot. A) Representative western blot. Aii) quantification of p-p65 activation for each stimulation condition. Data presented as mean ± SEM, N=3. B) Caco-2 cells were stimulated with 1 ng/mL or 1 µg/mL of LPS for 0, 1, 4, 8 or 24 h and cell viability was measured using Trypan blue. Conditions were run in duplicates and averaged. Data presented as mean ± SEM, N=3. Two-way ANOVA followed by Sidak’s multiple comparison post-hoc test. Ns; not significant. Differentiated Caco-2 cells were stimulated with 1 µg/mL of LPS for 0, 1, 4, 8 and 24 h and C) changes in TEER readings and D) permeability were measured using FITC Dextran. Data presented as mean ±SEM, N=3. One-way ANOVA followed by Dunnett’s multiple comparison post-hoc test, **P<0.005, ***P<0.0005, ****P<0.0001. E) Representative confocal images of Claudin-1 in Differentiated Caco-2 cells treated with 1 µg/mL of LPS for 0, 1, 4, 8 or 24 H. Damage indicated by white arrows. Scale bar = 10 µM.

| Target | Forward sequence | Reverse sequence |
|---|---|---|
| HCAR1 | CACCAGGAGGGAGGACAAA | CGTAGAGGAGCGATTGGGTC |
| TLR 2 | TGCAAGCAGGATCCAAAGGA | CCAGTGCTTCAACCCACAAC |
| TLR 4 | ACAACCTCCCCTTCTCAACC | TTGTCTGGATTTCACACCTGGAT |
| Ezrin | TGCTGCTGGATAGTCGTGTT | GCCGATAGTCTTTACCACCTGA |
| IAP | AGTTATCCTGCTCCCCACCT | TAGGAGGTGAAGGTCCAACG |
| SLC262A | TCCAACATTTCCCAGTTCCACA | TGTCCAGCTACAAGCTCTGC |
| CA2 | GACCCCTGGATGGCACTTAC | GTTTAGCGCTGCCAACCTTC |
| FABP1 | TATGGGCAGAGGTGGGAGTT | CAAGCCCCCAGTGAGTTTCT |
| FABP5 | AGCAGCTGGAAGGAAGATGG | CATTGCGCCCATTTTTCGCA |
| RBP2 | TCCTTGCCATCCACCACAAA | CCAGGTTCCATTCTGGTCCC |
| MCT1 | GTGTTCCTCTGTACTCTGGCC | ATTAGAAAGCTTCCTCTCCA |
| SMCT1 | AGCCACCATGATCCCAACTT | CTTTCCCCTCAAAGGGTGGTC |
| SMCT2 | AGATCCTCAGTGGCTGATGC | AGCAGAGAGGCTTTGCAGAG |
| GAPDH | CTTTTGCGTCGCCAG | TTGATGGCAACAATATCCAC |

**Supplemental Table 1: qPCR primer sequences.**
